# Single-cell transcriptomic analysis of human pleura reveals stromal heterogeneity and informs in vitro models of mesothelioma

**DOI:** 10.1101/2022.12.03.518966

**Authors:** Joanna Obacz, Reshma Nibhani, Taylor S. Adams, Jonas C. Schupp, Jose Antonio Valer, Niki Veale, Giuseppe Aresu, Aman S. Coonar, Adam Peryt, Giulia Biffi, Naftali Kaminski, Hayley Francies, Doris M. Rassl, Matthew J Garnett, Robert C. Rintoul, Stefan J. Marciniak

**Author notes:** Correspondence to: Stefan J. Marciniak, Cambridge Institute for Medical Research (CIMR), University of Cambridge School of Clinical Medicine, The Keith Peters Building, Cambridge Biomedical Campus, Hills Rd. Cambridge, CB2 0XY. Joint senior (Matthew J Garnett, Robert C. Rintoul, Stefan J. Marciniak).

## Abstract

The pleural lining of the thorax regulates local immunity, inflammation and repair. A variety of conditions, both benign and malignant including pleural mesothelioma, can affect this tissue. A lack of knowledge concerning the mesothelial and stromal cells comprising the pleura has hampered the development of targeted therapies. Here we present the first comprehensive single cell transcriptomic atlas of the human parietal pleura and demonstrate its utility in elucidating pleural biology. We confirm the presence of known universal fibroblasts and describe novel, potentially pleural-specific, fibroblast subtypes. We also present transcriptomic characterisation of multiple *in vitro* models of benign and malignant mesothelial cells, and characterise these through comparison with *in vivo* transcriptomic data. While bulk pleural transcriptomes have been reported previously, this is the first study to provide resolution at single cell level. We expect our pleural cell atlas will prove invaluable to those studying pleural biology and disease. For example, it has already enabled us to shed light on the transdifferentiation of mesothelial cells allowing us to develop a simple method for prolonging mesothelial cell differentiation in vitro.

## INTRODUCTION

The pleura regulates local immunity and wound-healing^1^. Perturbing these functions induces fibrosis and in the case of exposure to asbestos, chronic irritation can cause pleural mesothelioma, a rapidly progressive incurable cancer^1,2^. The lack of tractable *in vitro* models represents a significant barrier to the study of pleural biology. Although protocols for mesothelial cell extraction have been published^3,4^, *in vitro* culture is plagued by rapid transdifferentiation to fibroblast-like cells, the precise nature of which is unclear.

Epithelioid mesothelioma is the most differentiated primary pleural cancer and has a median survival of only 18 months^5^. This contrasts with the least differentiated subtype, sarcomatoid mesothelioma, which is typically fatal in under a year. Patients with histological evidence of both are said to have biphasic disease, which carries an intermediate prognosis. Although mesothelioma has a relatively modest mutational burden, BRCA1-associated protein 1 (*BAP1)*, merlin (*NF2*) and p53 (*TP53*) mutations are detected in 49%, 44% and 10% of case respectively^6^. The observation that *BAP1* is more commonly mutated in epithelioid than sarcomatoid tumours argues against a simple progression from epithelioid to sarcomatoid disease, and it remains uncertain if these subtypes even share the same cell of origin. Mesothelioma is a stromal-rich malignancy with cancer cells often representing only 5-25% of tumour nuclei^7^. Which cells are reprogrammed or recruited to become mesothelioma cancer-associated fibroblasts (CAF) is unknown. While the heterogeneity of tissue-specific fibroblasts has been well-studied in the lung^8^, skin^9^, heart^10^ and colon^11^, the fibroblasts of the pleura remain mysterious. It is known, however, that in some circumstances mesothelial cells can transdifferentiate to CAFs, since the peritoneal mesothelium contributes CAFs to pancreatic cancer and peritoneal metastasis^12,13^. In this regard, it would be valuable to understand better the relationships between mesothelial cells and their neighbouring fibroblasts.

Tissue fibroblasts have traditionally been defined by their spindle shape, the expression of canonical markers such as thymocyte differentiation antigen 1 (*THY1*) and platelet-derived growth factor receptor-α (*PDGFRA*), and as the source of extracellular matrix (ECM)^14^. Recently, it has become apparent that considerable diversity exists among fibroblasts with some possessing tissue-specific characteristics^15,16^. Analysis of single cell transcriptomes from various tissues suggests that both universal and specialised subtypes of fibroblasts exist^16^. In most tissues, dermatopontin-positive (*DPT*+) universal fibroblasts comprise subtypes marked by expression of peptidase inhibitor 16 (*PI16*) or collagen XV alpha chain 1 (*COL15A1*), with *PI16*+ cells expressing genes signifying an adventitial role in vascular niches, while *COL15A1*+ cells appear involved in secreting components of the basement membrane^15,16^. Lineage inference suggests that *PI16*+ cells possess the capacity to acquire the *COL15A1*+ state and may eventually develop into tissue-specific specialised fibroblasts, an example being the lipofibroblast in mature lung^17^. The heterogeneity of fibroblasts in pleura, however, has yet to be studied.

In addition to ECM, fibroblasts secrete inflammatory mediators and growth factors. This is relevant, for example, in cancers where CAFs modify prognosis and responses to therapy^18-20^. Alpha smooth muscle actin-expressing (*ACTA2*) myofibroblastic CAFs (myCAFs) play a major part in ECM deposition and TGFβ secretion, while *ACTA2*-low inflammatory CAFs (iCAFs) contribute to an immunosuppressive tumour microenvironment by secreting ligands such as CXCL1 and CXCL12^15^. Also, mesothelial-derived CAFs have been shown to modulate immunity through the expression of MHC class II (antigen presenting CAFs, apCAFs), perhaps to inhibit CD4+ T cell responses^12,21^. Much less is known about the role of fibroblasts and mesothelial cells in immune regulation in non-cancerous tissues.

To address the paucity of information available about the stromal cells of the pleura, we generated a single cell transcriptome atlas of healthy adult human parietal pleura, elucidating its cellular composition and phenotypic diversity. In doing so, we identified universal and novel populations of fibroblasts. Transcriptomic and functional studies demonstrated a key role for TGFβ in mesothelial to mesenchymal transition. Finally, by using the atlas to validate the transcriptomes of 2D mesothelial cell cultures, we were able to compare gene expression in healthy mesothelium with a panel of early passage mesothelioma 2D models revealing new phenotypic groups.

## RESULTS

### Mapping stromal heterogeneity in the human pleura

We isolated primary human parietal pleura from eight adult individuals undergoing video assisted thoracoscopic surgery for spontaneous pneumothorax (Table 1, Figure 1A). Tissue was disaggregated enzymatically with collagenase to a single-cell suspension and viable cells were enriched by fluorescence-activated cell sorting (FACS) with 7-aminoactinomycin D (7-AAD) staining. Single-cell RNA sequencing (scRNA-seq) was performed immediately using a droplet-based approach and data were combined to generate a pleural single-cell atlas of 64,514 cells (Figure 1B). Ten clusters were identified in most cases representing all participants (Figure 1B and Supplementary Figure S1). Based on the expression of canonical markers, major clusters were found to correspond to stromal, endothelial, immune, and mesothelial cells (Figure 1C-D and Supplementary Figure S2). To characterise the stromal and mesothelial components, we selected for clusters expressing *MSLN, PDGFRA, PDGFRB, LUM, ACTA2*, and *MYH11* and identified 32,750 cells (50.8% of cells sorted). Signature genes were cross-referenced to known markers of cell populations thereby identifying subclusters corresponding to mesothelial cells (n=11,535), smooth muscle cells (n=4245), pericytes (n=1194) and at least 5 populations of fibroblasts (Figure 1E-G and Supplementary Figure S3).

**Table 1.** Clinical characteristics of donors.

| Sample | Age (years) | Sex | Co-morbidities | Duration since most recent pneumothorax (days) | Total number of ipsilateral pneumothoraces |
| --- | --- | --- | --- | --- | --- |
| S1 | 16 | M | - | 11 | 1 |
| S2 | 27 | F | Autoimmune hepatitis<br>Hypermobility | 81 | 1 |
| S3 | 50 | M | - | 15 | 1 |
| S4 | 40 | M | - | 104 | 2 |
| S5 | 28 | M | - | 30 | 3 |
| S6 | 29 | M | Asthma | 300 | 1 |
| S7 | 17 | F | - | 25 | 2 |
| S8* | 34 | M | - | 4380 | 2 |
| N1 | 37 | M | - | 4 | 1 |
| N2 | 35 | F | Asthma | 8 | 1 |
| N3 | 29 | F | AD PCKD | 122 | 2 |
| N4* | 34 | M | - | 4380 | 2 |
| N5 | 54 | M | Sarcoidosis<br>Emphysema | 8 | 1 |

\* Donor S8 and N4 are the same individual
AD PCKD = autosomal dominant polycystic kidney disease

**Figure 1.**
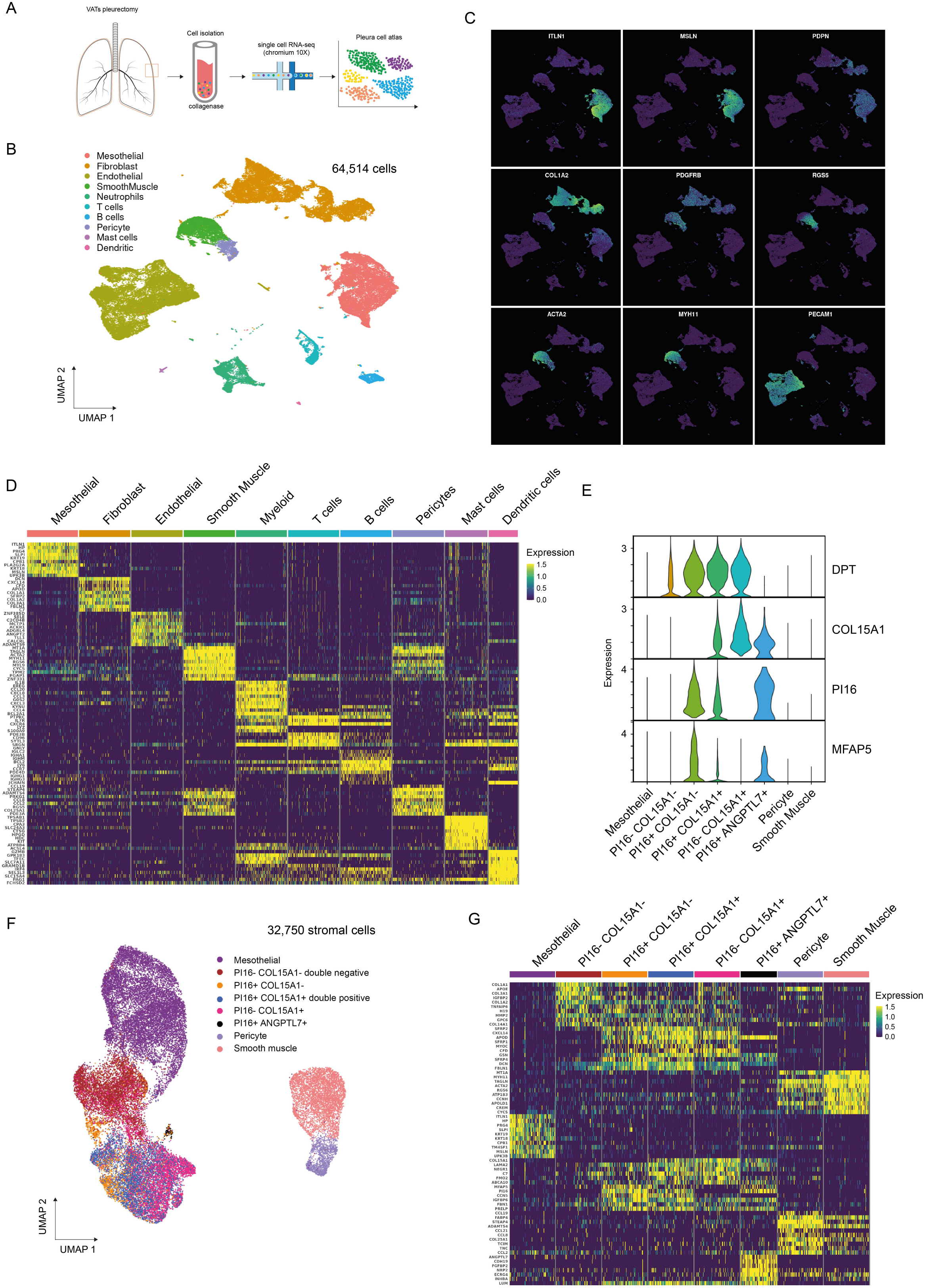
A single cell RNA sequencing (scRNA-seq) atlas of human parietal pleura. **A**. Study pipeline: samples obtained by video-assisted thoracic surgery were disaggregated to single cells with collagenase. Viable cells were collected by FACS and immediately processed by 10x Chromium Controller. **B**. Uniform manifold approximation and projection (UMAP) of 64,514 pleural cells from 8 donors are colour-coded by cell type (left). **C**. UMAP projection of canonical marker gene expression: mesothelial cells (ITLN1 interleukin 1; MSLN, mesothelin; PDPN podoplanin); fibroblasts (COL1A2 collagen type I alpha 2 chain; PDGFRB platelet derived growth factor receptor beta); pericytes (RGS5 Regulator Of G Protein Signaling 5); smooth muscle cells (ACTA2, smooth muscle α-actin-2; MYH11, myosin heavy chain 11); endothelial cells (PECAM1, platelet and endothelial cell adhesion molecule 1). **D**. Heat map of the relative average expression of the most strongly enriched genes for each cluster [log(fold change) of one cluster versus all others], grouped by cell type. All gene expression values are normalised across rows. **E**. Violin plots of DPT, COL15A1, PI16, MFAP5 expression. **F**. UMAP visualisation of 32,750 stromal cells. Eight populations identified through graph-based clustering are indicated by colour and annotated as Mesothelial, *PI16-COL15A1-, PI16+ COL15A1-, PI16+ COL15A1+, PI16-COL15A1+, PI16+ ANGPTL7+*, Pericyte and Smooth muscle. **G**. Heat map of the relative average expression of the most strongly enriched genes for each cluster in (F) [log(fold change) of one cluster versus all others], grouped by cell type. All gene expression values are normalised across rows.

To unpick the heterogeneity of fibroblasts in parietal pleura, we examined the expression of marker genes for known fibroblast subtypes. We observed *DPT*+ fibroblast populations that were also positive for either or both *PI16* and *COL15A1* (Figure 1E-F). *PI16* expression has been reported for fibroblasts in vascular niches^16^, while microfibril associated protein 5 (*MFAP5*) is detected in vascular adventitial fibroblasts of healthy distal lung^22^. In keeping with a putative vascular adventitial phenotype, *MFAP5* expression was detected in many of the pleural *PI16*+ *COL15A1-* fibroblasts (n=1864, Figure 1E-G). *PI16*-*COL15A1+* cells have been proposed to represent universal parenchymal fibroblasts with the ability to secrete basement membranes components^17^. We found these were abundant in human parietal pleura (n=4787, Figure 1E-G). We detected many pleural fibroblasts with an intermediate double-positive phenotype (n=4410). These *PI16*+ *COL15A1+* cells expressed genes characteristic of both *PI16*+ *COL15A1-* fibroblasts *(CCN5, IGFBP6, FBN1, PRELP*) and *PI16*-*COL15A1+* fibroblasts (*LAMA2, NEGR1, C7FMO2, ABCA10*), although unlike *PI16*+ *COL15A1-* cells, they expressed relatively low levels of MFAP5 (Fig 1E-G and Supplementary Figure S4). In addition, a rare subpopulation of *PI16+* fibroblasts (n=80 cells, 0.2% of stromal cells) was detected, which selectively expressed angiopoietin-related protein 7 (*ANGPTL7*), a gene recently implicated in regulating stemness and expressed by resting hair follicle stem cells^23,24^ (Fig 1E-G and Supplement Figure S5).

### Distinct functional subsets of fibroblasts are detected in benign human pleura

Gene set enrichment analysis of the stromal populations was performed to gain functional insight into the subpopulations. In all *PI16+* fibroblast clusters, pathway analysis of differentially expressed genes identified patterns associated with focal adhesion, ECM-receptor interaction, PI3K-Akt signalling and TGFβ signalling (Figure 2A). Similar pathways were detected in *PI16-COL15A1*+ fibroblasts, although TGFβ signalling was less prominent. A substantial population of fibroblasts failed to express either *PI16* or *COL15A1* (n=4635, Figure 1F), and these double-negative cells lacked evidence of PI3K-Akt and TGFβ signalling, but expressed genes associated with ECM-interaction (Figure 2A). The *ANGPTL7*+ subpopulation of *PI16+* fibroblasts showed evidence of WNT and TGFβ signalling, with ECM-related functions being less evident (Figure 2A). This combination suggests a progenitor-like role^25-27^, as has been suggested for other *PI16*+ fibroblasts^16^, so it is noteworthy that other stemness genes (including *SOX8*^28^ and *SHISA* family members^29^) were selectively expressed in this population (Figure 2B and Supplementary Figure S6). A similar population of cells named endoneurial fibroblasts was previously described in peripheral nerves^30^.

**Figure 2.**
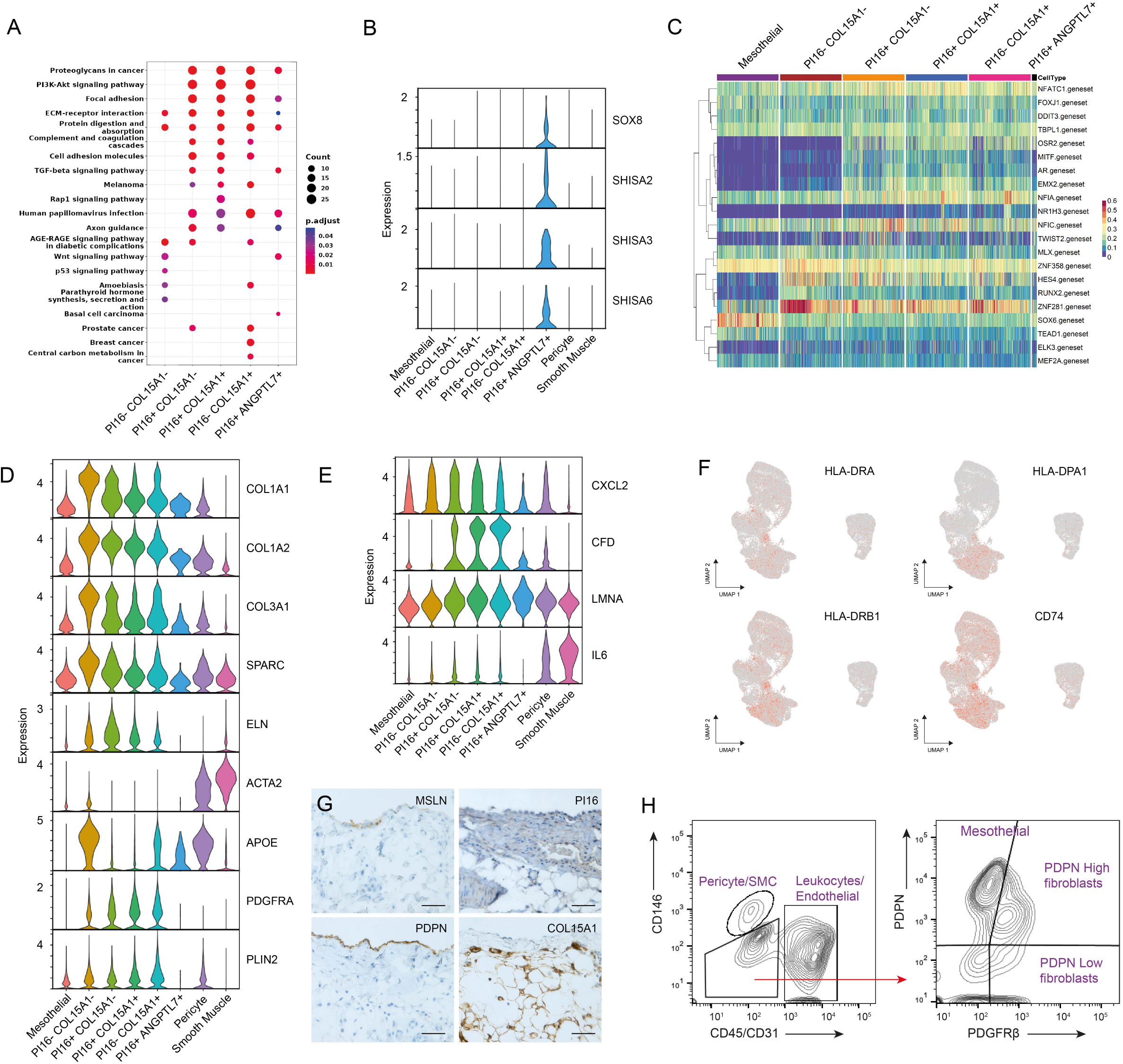
Stromal heterogeneity in healthy pleura. **A**. KEGG pathway gene set enrichment analysis of pleural fibroblasts. **B**. Violin plots of SOX8, SHISA2, SHISA3, SHISA6 expression. **C**. Heat map of Gene Regulatory Network (GRN) analysis. **D**. Violin plots of COL1A1, COL1A2, COL3A1, SPARC, ELN, ACTA2, APOE, PDGFRA, PLIN2 expression. **E**. Violin plots of CXCL2, CFD, LMNA, IL6 expression. **F**. UMAP visualisation HLA-DRA, HLA-DPA1, HLA-DRB1, CD74. **G**. Representative immunohistochemically stained images of mesothelin, podoplanin, COL15A1 and PI16 in human pleura tissue sections. Bar = 50 μm. **H**. Fluorescence-activated cell sorting (FACS) of pleural stromal components.

Gene regulatory network (GRN) analysis was performed to infer activity of transcription factor regulons across stromal populations. Mesothelial cells were found to display greater SOX6 regulon activity than was detected in pleural fibroblasts (Figure 2C). SOX6 has been shown to mark mesothelioma in pleura, peritoneum and tunica vaginalis, but has never previously been studied in benign pleura^31-33^. Consistent with the GRN analysis, our scRNA-seq atlas confirmed that SOX6 was more robustly expressed in mesothelial cells compared with most other stromal components of the pleura (Supplementary Figure S7). MITF, AR and EMX2 genesets were also detected in *PI16*+ cells as were the transcription factors themselves, although their functional significance remains unclear.

Double negative *PI16-COL15A1*-fibroblasts showed activation of the regulons of HES4, a Notch responsive transcription factor previously implicated in regulating stromal cell specialisation^34^, and ZNF281, an EMT-inducing transcription factor^35,36^ (Figure 2C). Both ZNF281 and HES4 regulons were active in other fibroblast populations but at lower levels. The transcription factors HES4 and, to a lesser extent, ZNF281 were expressed in a proportion of *PI16-COL15A1*-population (Supplementary Figure S7). *PI16-COL15A-* cells strongly expressed components of the collagen-rich ECM (*COL1A1, COL1A2, COL3A1* and *SPARC)*, although other canonical myofibroblast markers (*ELN, ACTA2, TAGLN MYH11)* were detectable only at low levels (Figure 2D & Supplementary Figure S8)^8,16^. These double-negative cells also expressed high levels of *APOE*, a marker of lipofibroblasts, but relatively low levels of other lipofibroblast genes (*PDGFRA* and *PLIN2*) (Figure 2D). *PI16-COL15A-* double-negative pleural fibroblasts therefore display a phenotype intermediate between lipofibroblasts and myofibroblasts.

A role has emerged for fibroblasts in local immune regulation^15,21^. We therefore searched for evidence of immune modulatory fibroblasts in benign pleura. *DPT+* marks both universal fibroblasts and inflammatory fibroblasts (Figure 1)^15,16,21^, and additional inflammatory fibroblasts-associated genes (*CXCL12, CFD and LMNA*) were detected in some *PI16*+ and *COL15A1*+ double-positive fibroblasts, while other canonical iCAF-like marker genes (*IL6, AGTR1, HAS1, HAS2*) were less expressed (Figure 2E and Supplementary Figure S9) as likely they require additional tumour cues. In benign pleura, we detected low levels of HLA-DRA, HLA-DPA1 and HLA-DRB1 expression in some universal fibroblasts (Figure 2F). The expression of CD74, which encodes the invariant MHC class II gamma chain, suggests that functional MHC class II may be synthesised.

Immunohistochemical analysis confirmed mesothelin (MSLN) and podoplanin (PDPN) expression by the mesothelial monolayer (Figure 2G). To confirm the presence of the stromal cell populations detected by scRNA-seq, we used a flow cytometry panel comprising PDPN, PDGFRβ and CD146 to distinguish mesothelial cells, fibroblasts, pericytes and smooth muscle. Fresh tissue was dissociated with collagenase and single, viable cells were enriched. Immune cells and endothelial cells were depleted based on CD45 and CD31 expression respectively, while pericytes and smooth muscles were identified by CD146. CD45-CD31-CD146-triple-negative cells could be further separated into PDPN+ PDGFRβ-mesothelial cells, PDPN^low^ universal and PDPN^high^ fibroblasts (Figure 2H).

These data demonstrate that benign human pleura comprises multiple fibroblast populations including known *DPT*+ universal subtypes. An abundant pleural *PI16-COL15A1-* double-negative population was found to display features of lipo- and myofibroblasts, and may serve as a source of fibrillar collagens in the parietal pleura. A low abundance population of *PI16*+ *ANGPTL7+* cells expressed genes suggesting a progenitor-like behaviour potentially related to peripheral nerves. Some *DPT+* fibroblasts expressed genes involved in immune modulation, while certain mesothelial cells and *DPT*+ fibroblasts expressed MHC class II, which suggests a potential immunomodulatory role in normal pleura.

### TFGβ drives mesothelial-to-mesenchymal transition

Single cell data from fresh pleura revealed that several genes highly expressed in mesothelial cells (*KRT8, KRT18, PDPN*) were expressed at lower levels in *PI16-COL15A1-* double-negative fibroblasts (Figure 3A-B). Although, mesothelial-to-mesenchymal transition (MMT) has been studied in cancer, organ development and fibrosis^37^, little is known about MMT in non-cancerous pleura nor the cellular fate of transitioning mesothelial cells. To address this, we performed pseudo-time analysis using PHATE (Potential of Heat Diffusion for Affinity-based Trajectory Embedding), a bioinformatic tool enabling 2D visualisation of trajectory structure in high-dimensional data^38^. This suggested that mesothelial cells may transition preferentially to *PI16-COL15A1-* fibroblasts (Figure 3C). *PI16-COL15A1-* fibroblasts may also transition to PI16+ and/or COL15A1+ fibroblasts. In an effort to understand the drivers of pleural MMT, we modelled ligand-receptor interactions using the NicheNet algorithm hoping to identify ligands that promote fibroblast growth and survival^39^. Ligand activities were ranked and Pearson coefficients were calculated for the correlation between targets of a given ligand and the expression of genes in the receiver cell (Figure 3D). With mesothelial cells as senders and *PI16-COL15A1-* double negative fibroblasts as receivers, this analysis nominated TGFB1 as a putative ligand signalling from mesothelial to *PI16-COL15A1-* pleural fibroblasts.

**Figure 3.**
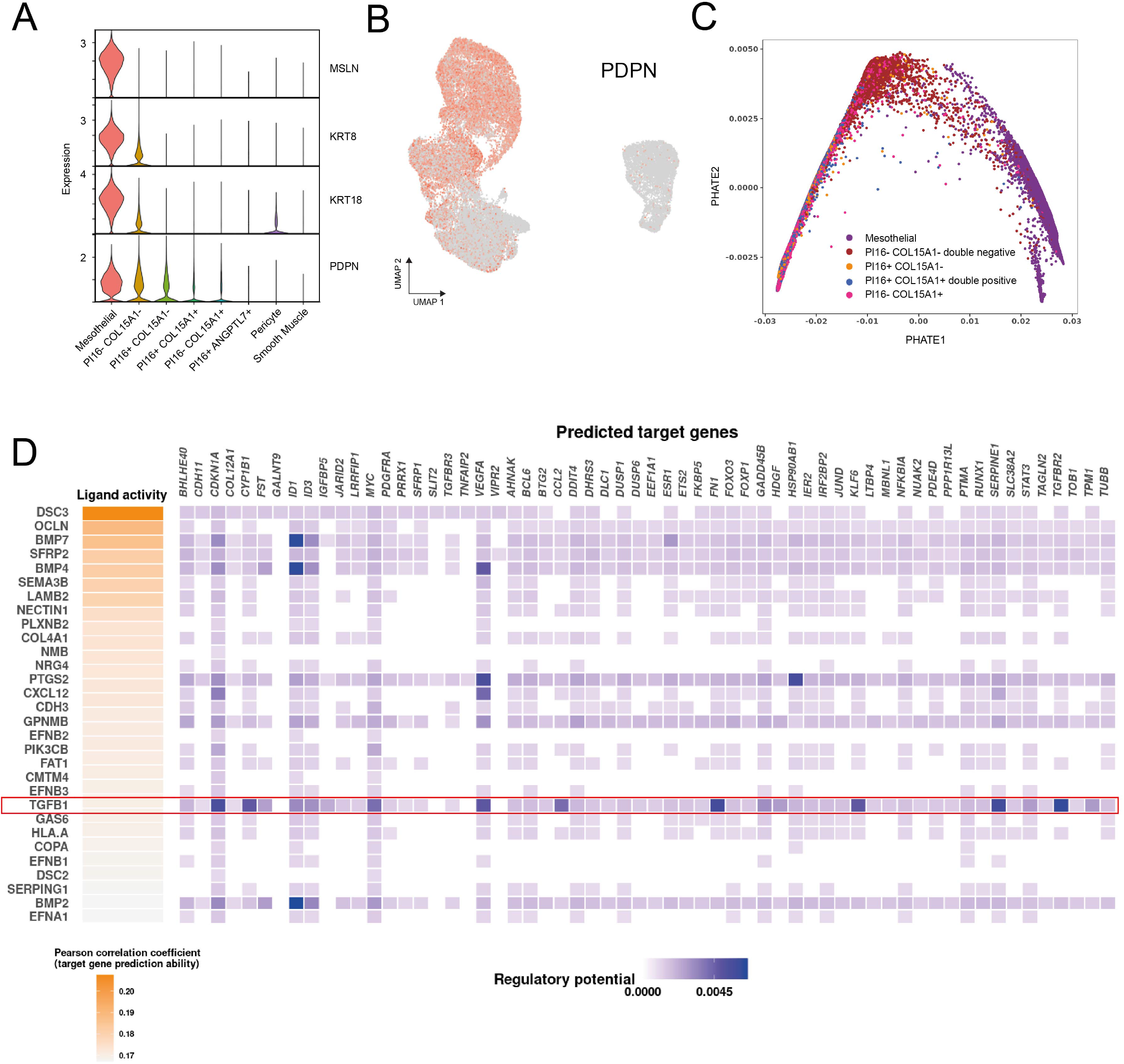
Isolation and dedifferentiation of parietal pleural mesothelial cells. **A**. Violin plots of KRT8, KRT18, PDPN expression. **B**. UMAP visualisation of APOE expression in mesothelial and stromal cells. **C**. Potential of heat diffusion for affinity-based transition embedding (PHATE) visualisation of pseudotime trajectory of mesothelial trans-differentiation into activated fibroblasts. **D**. Intracellular communication between mesothelial cells and activated fibroblasts as determined by NicheNet algorithm. Ligand activity prediction showing 20 mesothelial ligands with the highest likelihood (Pearson correlation coefficient) of affecting the gene expression in the receiver activated fibroblasts. Ligands specifically interacting with activated fibroblasts (but not universal fibroblasts) are depicted in red. Ligand-target matrix denoting the regulatory potential between mesothelial-ligands and target genes in activated fibroblast.

To examine MMT in vitro, we isolated primary mesothelial cells by limited trypsinisation of fresh pleura, which unlike collagenase treatment does not mobilise large numbers of fibroblasts (Figure 4A). Mesothelial cells from individuals underdoing pleurectomy for pneumothorax, formed monolayers with a cobblestone morphology and expressed MSLN and PDPN, which are well-validated markers of mesothelial cells (Table 1, Figure 4B-D). Electron microscopy confirmed the presences of abundant surface microvilli typical of this cell type (Figure 4B). Subsequently, during in vitro culture, primary mesothelial cells changed morphology, acquiring an elongated, spindle-shape resembling fibroblasts (Figure 4C). In parallel, MSLN staining weakened, though PDPN staining was retained for at least 5 days (Figure 4D).

**Figure 4.**
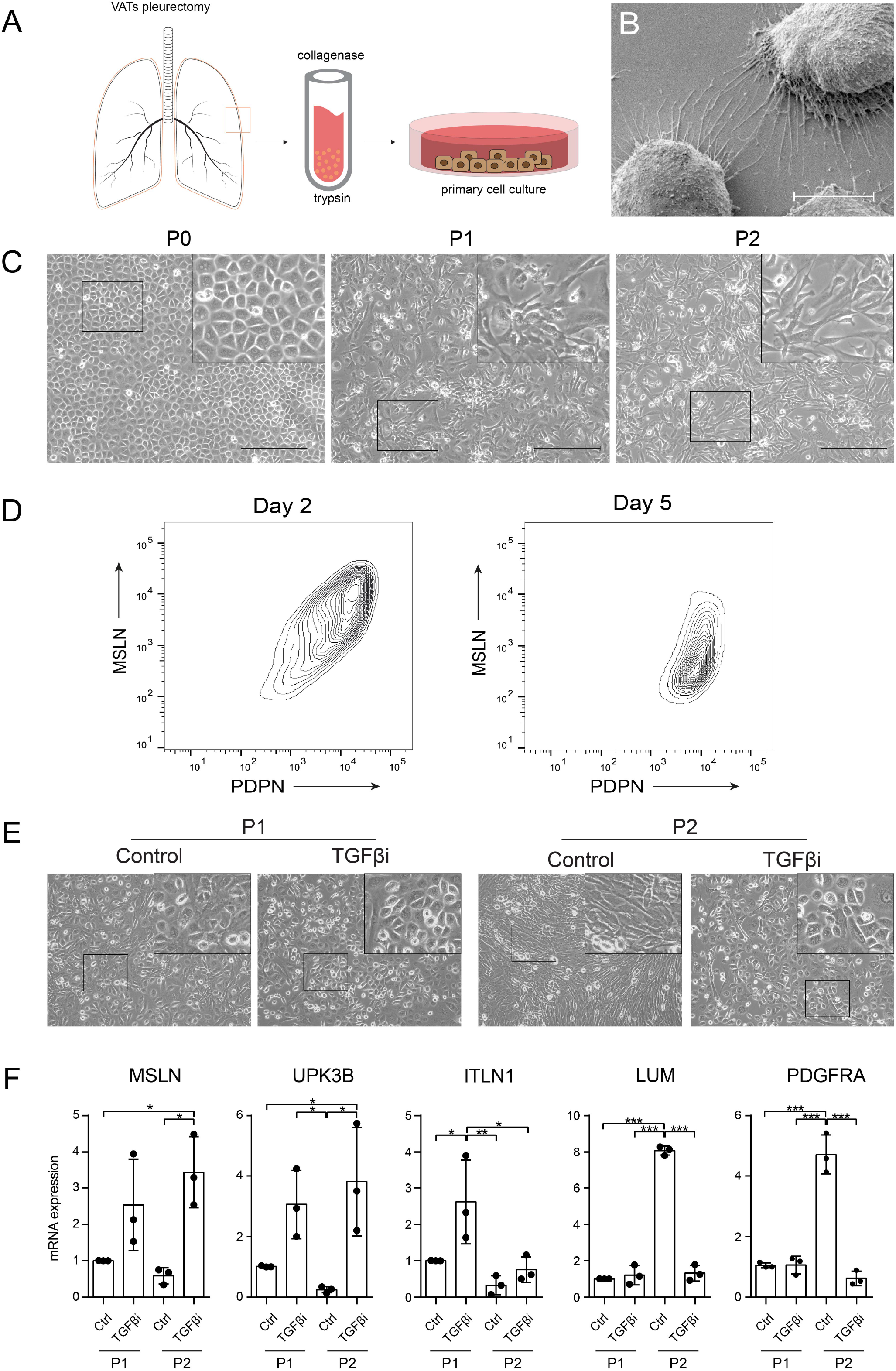
TGFβ signalling regulates pleural mesothelial-to-mesenchymal transition. **A**. Mesothelial cells isolation pipeline: samples obtained by video-assisted thoracic surgery were disaggregated to single cells with collagenase. Viable cells were collected by FACS and immediately processed by 10x Chromium Controller. **B**. Scanning electron micrograph of low confluence trypsin-isolated mesothelial cells. Scale bar = 20μm **C**. Phase contrast images of primary mesothelial cell cultures at passage 0 (P0), passage 1 day 7 post-surgery (P1) and passage 2, day 14 post-surgery (P2). Scale bar = 300μm. Insert shows zoomed image of boxed region. **D**. FACS of trypsin-isolated mesothelial cell cultures and day 2 and day 5, stained for MSLN and PDPN. **E**. Phase contrast images of primary mesothelial cell cultures at P1 and P2, without (Ctrl) or with TGFβ inhibitors SB431542 and Noggin (TGFβ inhibitors SB431542 and Noggin 10μM and 100ng/ml, respectively). Insert shows zoomed image of boxed region. **F**. Mesothelial and fibroblast gene expression with or without TGFβ inhibition qPCR. Three independent experiments are presented (dots for each experiment and histogram for mean). *p < 0.05, **p < 0.01, ***p < 0.001; one-way ANOVA with Tukey”s multiple comparisons test.

To test whether TGFβ sustains MMT in vitro, we cultured low-passage mesothelial cells from three healthy donors in the presence or absence of the TGFβ inhibitors SB431542 and Noggin. As expected, in the absence of TGFβ inhibition, mesothelial cells started to lose their cobblestone morphology on passaging (Figure 4E). This was associated with a reduction of mesothelial marker gene expression (*MSLN, UPK3B, ITLN1)* and a corresponding increase in fibroblast marker gene expression (*LUM, PDGFRA)* (Figure 4F). By contrast, cultures treated with the TGFβ inhibitors, retained mesothelial cell morphology and marker gene expression without increased fibroblast gene expression. This suggests that TGFβ, derived locally from mesothelial cells, may drive trans-differentiation of mesothelial cells to fibroblasts when grown in vitro and that this is susceptible to pharmacological modulation.

### Linking scRNA-seq atlas with 2D models of the mesothelium and mesothelioma

Analysis of *in vitro* models of mesothelioma has been limited by a paucity of authentic benign controls. To determine if early-passage primary mesothelial cells recapitulate their counterparts *in vivo*, we used bulk RNA-seq to examine the transcriptomes of early-passage cultures (5-6 days post-isolation) from 5 further individuals. In parallel, we performed bulk RNA-seq of a panel of 21 early passage cell models of mesothelioma (passage 5-8)^40,41^. The 1,000 most differentially expressed genes were selected for hierarchical clustering using a Pearson correlation coefficient as the distance metric and Ward”s linkage rule^42^ (Figure 5A). All 5 primary benign cultures (N1-N5) clustered together, as did those from mesothelioma. To compare the bulk RNA-seq data with our scRNA-seq atlas of parietal pleura we used Single-Cell Identification of Subpopulations with Bulk Sample Phenotype Correlation (SCISSOR) analysis^43^. This confirmed that our primary cultures from healthy individuals mapped well to cells in the mesothelial cluster (Figure 5B).

**Figure 5.**
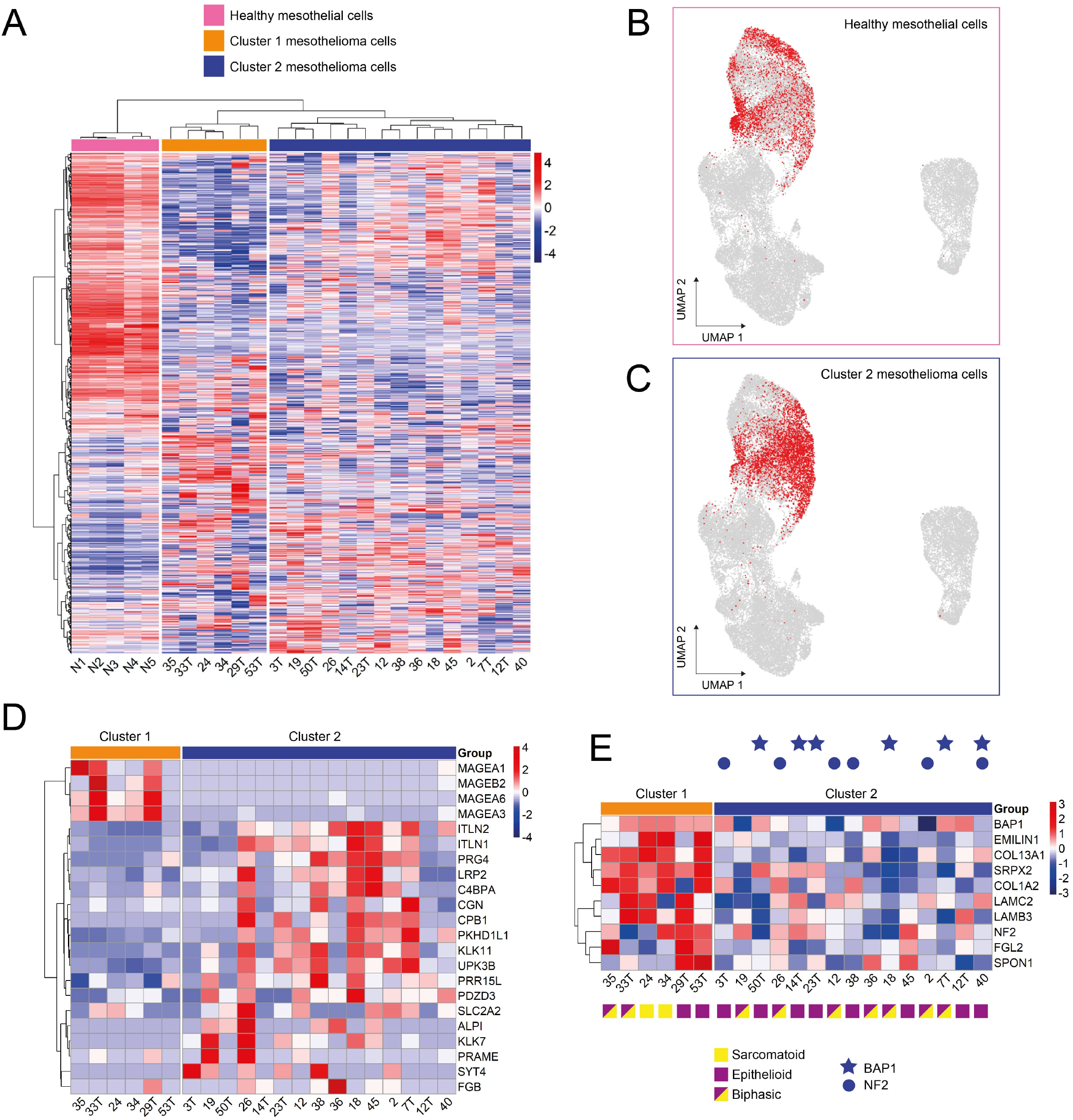
Characterisation of 2D models of mesothelioma. **A**. Unsupervised classification of gene expression data (Bulk RNA seq) of 5 primary mesothelial cell cultures (N1-5) and 21 mesothelioma cell lines. Count data transformation was performed by using the variance stabilizing transformation implemented in the DESeq2 package. For each gene, the dispersion was calculated to measure its variance among samples, and thus the 1,000 genes with highest dispersions were selected for clustering analysis. Hierarchical clustering analysis was performed using the Pearson correlation coefficient as the distance metric and Ward”s linkage rule. **B**. Single-Cell Identification of Subpopulations with Bulk Sample Phenotype Correlation (SCISSOR) analysis to identify stromal cells in the scRNA-seq atlas with similar gene expression to healthy mesothelial cell cultures. **C**. SCISSOR analysis to identify stromal cells in the scRNA-seq atlas with similar gene expression to Cluster 2 mesothelioma cell lines. **D**. Unsupervised classification of gene expression data (Bulk RNA seq) 21 mesothelioma cell lines divided into clusters 1 and 2 defined in “A”. Hierarchical clustering analysis was performed using the Pearson correlation coefficient as the distance metric and Ward”s linkage rule. Genes were selected from those most differentially expressed in epithelioid vs sarcomatoid mesothelioma^44^. **E**. Heat map of “ECM structural constituents” geneset (*EMILIN1, COL1A2, FGL2, LAMB3, COL13A1, LAMC2, SRPX2, SPON1)*) in cluster 1 compared to cluster 2. BAP1 mutations indicated by star. NF2 mutation indicated by circle. Cell lines derived from sarcomatoid = tumour yellow square, from epithelioid = purple square, from biphasic tumour = purple/yellow square.

Having generated a valid control dataset, we compared this with the transcriptomes of early passage mesothelioma. By Gene Ontology (GO) Term analysis, mesothelioma cells exhibited higher genes expression associated with growth factor activity and ECM production, but lower immune receptor activity (Supplementary Figure S10). By KEGG pathway analysis, mesothelioma cells had increased focal adhesion, ECM-receptor interaction, and PI3K-Akt signalling, but less gene expression associated with inflammation (Supplementary Figure S10). Within the 21 mesothelioma cell lines, two distinct clusters were observed, which could not be explained by systematic procedural differences in their derivation^40,41^. When these clusters were examined by SCISSOR (Figure 5C, compare with Figure 1F), cluster 2 showed similarity to mesothelial cells *in vivo*, while cluster 1 was dissimilar to all healthy pleural cell populations. Cluster 2 was enriched for genes observed to be expressed preferentially in epithelioid mesotheliomas compared with sarcomatoid disease: *ITLN1, ITLN2, PRG4, LRP2, C4BPA, CGN, CPB1, PKHD1L1, KLK11* and *UPK3B* (Figure 5D)^44^. Conversely, several genes reported to be expressed in sarcomatoid mesothelioma were preferentially expressed in cluster 1, including *MAGEA1, MAGEA3, MAGEB2, and MAGEA6*.

We next compared cluster 1 (sarcomatoid-like) with cluster 2 (epithelioid-like). Consistent with a sarcomatoid-like phenotype, by GO Term analysis cluster 1 showed a significant upregulation of the ECM structural constituents geneset: *EMILIN1, COL1A2, FGL2, LAMB3, COL13A1, LAMC2, SRPX2, SPON1* (p = 0.01; Figure 5E). Genomic sequencing further supported the notion that cluster 1 might represent the sarcomatoid phenotype while cluster 2 represents epithelioid mesothelioma, since *BAP1* and *NF2* pathogenic variants, which are more common epithelioid tumours^2^, were detected only in cluster 2. Interesting, of the 21 early passage mesothelioma cultures examined, only two had originated from individuals with proven sarcomatoid mesothelioma, both of these were in cluster 1.

These data suggest that early passage parietal mesothelial cells isolated by limited trypsinisation efficiently model healthy mesothelial cells in vivo. This provides a valuable control for comparison with disease states such as malignancy, while TGFβ inhibition may prove useful in allowing healthy mesothelial cell cultures to be maintained in vitro for longer periods to facilitate such work. Our comparison highlights changes in mesothelial factors that influence their local environment including the extra-cellular matrix and communication with the immune system. We also showed that in vitro, early passage mesothelioma cell cultures adopt two main phenotypes that show similarity with epithelioid and sarcomatoid mesothelioma respectively, but this does not always reflect the diagnostic label of the donor.

## DISCUSSION

We analysed 64,514 cells by scRNA-seq from eight individuals to define the cellular components of the human parietal pleura. In doing so, we have identified known and previously undescribed fibroblast subtypes. Comparison of the *in vivo* transcriptomes of parietal mesothelial cells with those of healthy mesothelial cells grown in 2D culture, validated our protocol for mesothelial cell isolation, while comparisons with early passage mesothelioma cells revealed them to partition between two phenotypic groups, which may have value in identifying subtype-specific therapies.

Single-cell RNA-sequencing is revolutionising our understanding of complex biological systems in heath and disease. Although the physiological functions of human pleura are well described^1^, the interactions between pleural cell components have yet to be studied in detail. Since these interactions are fundament to the aetiology of pleural pathologies, including benign pleural thickening and malignant pleural mesothelioma, we set out to generate a parietal pleural cell atlas. In addition to mesothelial cells, pericytes and smooth muscles, we uncovered the functional heterogeneity of parietal pleural fibroblasts. Consistent with recent work^16^, we identified *COL15A1+* and *PI16+* universal fibroblasts corresponding to parenchymal and adventitial populations respectively. A *COL15A1-PI16-* subtype lacking universal fibroblast markers was found to be abundant in the parietal pleural. This subtype displayed a phenotype related to lipofibroblasts, for example positivity for APOE, while simultaneously showing myofibroblast features, including expression of collagen-rich ECM component *COL1A1, COL1A2, COL3A1* and *SPARC*. The latter suggests a potential role as the pleural source of fibrillar collagens, but the relevance of lipofibroblast features remains to be explored. We uncovered a rare population of *PI16+ ANGPTL7*+ fibroblasts, whose transcriptomic signature suggests a potential progenitor-like function. Similar cells named endoneurial fibroblasts have been described in peripheral nerves and so these may derive from innervation of the pleura^30^. It is tempting, however, to speculate that these may also be relevant to known pleural responses to chronic local inflammation, namely extrapleural fat deposition, fibrosis and calcification.

Mesothelial cells are highly versatile and under appropriate conditions can transdifferentiate into adipocytes or osteoblasts^45^, myofibroblasts^46^ and even vascular smooth muscle cells^47^. Here, using pseudotime modelling and *in vitro* assays, we further support the multipotent nature of pleural mesothelial cells with evidence for transdifferentiation to fibroblasts under physiological conditions. In pulmonary fibrosis, TGFβ has a role in epithelial-to-mesenchymal and mesothelial-to-mesenchymal transition^48,49^. Here, we demonstrate that antagonising TGFβ signalling delays transdifferentiation of mesothelial cells in culture. Conversely, the networks maintaining mesothelial differentiation remain unclear. We observed that SOX6 regulon activity is more readily detectable in mesothelial cells than in other stromal components, while SOX6 itself is detectable predominantly in mesothelial cells. It has recently been noted that SOX6 is a marker of mesothelioma, but to our knowledge, ours is the first description of SOX6 activity in benign pleura^31-33^. Some expression of SOX6 was detected in *COL15A1-PI16-* double-negative pleural fibroblasts, although without obvious activation of its regulon above that seen in other fibroblasts. By contrast, double-negative fibroblasts showed expression of HES4 and activation of its regulon, both of which were absent in mesothelial cells. Since HES4 is a Notch-responsive transcription factor and has been implicated in stromal cell specialisation^34^, further work will be required to clarify its role in pleural stromal function.

We observed that some *PI16*+ and *COL15A1*+ double-positive pleural fibroblasts express genes previously reported to be associated with an inflammatory fibroblast phenotype (*CXCL12, CFD and LMNA*), although additional cues may be necessary to manifest a full iCAF-like phenotype, since other canonical iCAF marker genes were not expressed by this subpopulation (e.g. *IL6, AGTR1, HAS1, HAS2*)^21^. apCAFs are another type of immune-modulating fibroblasts, which in human and murine pancreatic adenocarcinoma express MHC class II and modulate CD4+ T cells^12,21^. Indeed, it has been suggested that peritoneal mesothelial cells may be progenitors of apCAFs in pancreatic cancer^12^. In this regard it is intriguing that detectable HLA-DRA, HLA-DPA1 and HLA-DRB1 expression was observed in some universal fibroblasts.

In the present work, we provided a simple protocol for the establishment of *in vitro* benign pleural mesothelial cell cultures whose transcriptomes mirror those of parietal mesothelial cells *in vivo*. To showcase the utility of our pleural scRNA-seq data, we then explored the transcriptome of primary low passage mesothelioma cells lines. Unsupervised clustering of these malignant cells based on their transcriptomes yielded two groups: one sharing marked similarity with healthy mesothelial cells *in vivo*, while the second was dissimilar to any healthy pleural cell type, including mesothelial cells and fibroblasts. Comparison with published tumour sequencing data, suggests the cluster 1 (the mesothelial-like group) may serve as a model for epithelioid mesothelioma, while the cluster 2 cell lines more closely resemble sarcomatoid mesothelioma. This information will help guide subtype-targeted drug discover efforts.

In summary, we have generated a single cell atlas of human parietal pleural revealing heterogeneity among the stromal component including the identification of novel fibroblast subtypes. We have shown utility of these data by identifying and manipulating signals regulating the transdifferentiaton of mesothelial cells *in vitro* and have used this atlas to validate in vitro models of mesothelioma.

## MATERIALS AND METHODS

### Patients

This study was carried out under the Royal Papworth Hospital Research Tissue Bank Generic Research Ethic Committee approval (Tissue Bank project number T02537). Patients who underwent the video-assisted thoracoscopic surgery for pneumothorax in Royal Papworth Hospital in Cambridge, UK between 2019 and 2022 were enrolled in this study (Table 1). All the patients provided written informed consent for sample collection and data analyses prior to surgery. All samples were provided as linked-anonymised, with the linkage key held by the Tissue Bank.

### Tissue processing for mesothelial cells isolation

Immediately after the surgery, pleura specimens (typically 3-5 cm^3^) were immersed in the cold RPMI media supplemented with penicillin (100 units/mL) and streptomycin (100 μg/mL) (Gibco). Mesothelial cells were isolated by incubating small fragments of pleura (∼0.5cm^3^) in 0.25% Trypsin/0.01% EDTA at 37°C with 150 rpm agitation for 30 mins. Following that, detached cells were collected by centrifugation at 800g for 2 mins and red blood cells were removed by incubating the pellet with 1× red blood cell lysis buffer (Life Technologies) for 7 mins in RT. After washing with PBS, mesothelial cells were pelleted at 800g for 2 mins and resuspended in 50:50 v/v of RPMI (supplemented with 10% FBS, 1× penicillin/streptomycin, 2 μg/ml heparin (Sigma) and 1 μg/ml hydrocortisone (Sigma)) and BEBM™ Bronchial Epithelial Cell Growth Basal Medium supplemented with the BEGM™ Bronchial Epithelial Cell Growth Medium Single Quots™ Supplements and Growth Factors (Lonza). Fresh media were added to the cells every 2-3 days until the confluent monolayer of mesothelial cells was obtained (typically between 5-10 days post-isolation).

### Primary mesothelioma cell culture

All cell lines were obtained from Mesobank UK^40^ and were cultured in RPMI-1640 growth media supplemented with l-glutamine (2 mM), penicillin (100 U/mL), streptomycin (100 *μ*g/mL), hEGF (20 ng/mL), hydrocortisone (1 *μ*g/mL), heparin (2 *μ*g/mL) and 10% FBS at 37 °C and 5% CO_2_ as previously described^41^.

### Tissue processing for single cell RNA sequencing (scRNA-seq)

Immediately after surgical resection, pleura specimens were immersed in the cold RPMI media supplemented with 100 units/mL of penicillin and 100 μg/mL of streptomycin (Gibco) and, whenever possible, processed within 1 hr of receipt. For the overnight storage at 4°C, tissue was transferred into 10 mL cold HypoThermosol FRS solution (Sigma) to stabilize the intact tissue for scRNA-seq^50^. Following that, pleura was washed three times in PBS, minced with scalpels and enzymatically digested in RPMI supplemented with collagenase I (Life Technologies, 5 mg/ml) and DNase I (Sigma, 100 μg/ml) for 2.5 hrs at 37°C with 150 rpm agitation. Digested tissue was filtered through a Falcon® 100μm cell strainer (Corning), washed with PBS and centrifuged at 800g for 2 mins. Cell pellet was resuspended in 1× red blood cell lysis buffer (Life Technologies) and incubated for 7 mins in RT. Following dilution, pelleting, and resuspension in RPMI supplemented with 2% FBS cells were counted, and viability was determined using a Countless Automated Cell Counter Hemocytometer (Invitrogen) and trypan blue. To generate a single cell suspension, cells were stained with 7-AAD (Thermofisher), and live cells were collected using BD FACSMelody™ Cell Sorter (BD Biosciences). After sorting, cells were pelleted, resuspended in RPMI supplemented with 2% FBS and counted using trypan blue and hemocytometer. A total of 16,000 cells were submitted to 10x Genomics and immediately loaded on 10x 3” chip.

### Single cell RNA-seq library preparation, next-generation sequencing and data processing

scRNA-seq libraries were generated using the 10x Chromium platform 3′ v3.1 kit (10x Genomics) following the manufacturer”s recommendations and targeting 10,000 cells per sample. Library quality was assessed following cDNA synthesis and after completion of the 3” Gene Expression Libraries on the Agilent TapeStation system employing a DNA High Sensitivity D5000 and D1000 chips, respectively (all Agilent). Single cell libraries were sequenced on Illumina NovaSeq 6000 instrument using read 1: 28bp and read 2: 90bp length sequencing and aiming at 20,000 reads/cell. Raw reads were processed using the Cell Ranger 2.1.0 pipeline with default and recommended parameters. Gene-Barcode matrices were generated for each individual sample by counting unique molecular identifiers (UMIs) and filtering non-cell associated barcodes. Finally, we generated the gene-barcode matrix containing the barcoded cells and gene expression counts. This input was then imported into the Seurat (v4) R toolkit for quality control and downstream analysis. All functions were run with default parameters, unless specified otherwise. Low quality cells (<200 genes/cell, <3 cells/gene and >20% mitochondrial genes) were excluded.

### Identification of cell types and subtypes by Uniformed Manifold Approximation and Projection (UMAP)

The Seurat package implemented in R was applied to identified major cell types. Highly variable genes were generated and used to perform PCA. Significant principal components were determined using JackStraw analysis and visualisation of heatmaps focusing on PCs 1 to 20. PCs 1 to 10 were used for graph-based clustering (at res = 0.8) for all samples. These groups were projected onto UMAP analysis run using previously computed principal components 1 to 10.

### Cluster markers identification and downstream analyses

The cluster specific marker genes were identified by running the FindAllMarkers function in Seurat package to the normalised gene expression data. We used clusterProfiler Bioconductor package to perform biological processes enrichment analyses, PySCENIC for gene regulatory network interference and NicheNet^39^ for ligand-receptor interaction analysis.

### Unsupervised classification of bulk RNA-seq gene expression data

Count data transformation was performed using the variance stabilising transformation implemented in the DESeq2 package. For each gene, the dispersion was calculated to measure its variance among samples, and thus the 1,000 genes with highest dispersions were selected for clustering analysis. Hierarchical clustering analysis was performed using the Pearson correlation coefficient as the distance metric and Ward”s linkage rule^42^.

#### Gene regulatory network (GRN) analysis

The pySCENIC package (version 0.11.2), a Python-based implementation of the SCENIC pipeline, with two gene-motif rankings (hg38 refseq-r80 10kb_up_and_down_tss and hg38 refseq-r80 500bp_up_and_100bp_down_tss) from the RcisTarget database was used to investigate the gene regulatory network of transcription factors (TFs)^51^. Areas under the recover curve (AUC) were calculated with a threshold set at 0.35.

### Integration of phenotype-associated bulk expression data and single-cell data

Bulk RNA-seq and scRNA-seq data were integrated using Single-Cell Identification of Subpopulations with Bulk Sample Phenotype Correlation (SCISSOR) analysis^43^. https://github.com/sunduanchen/Scissor/

### Flow cytometry analysis

Pleura specimens were minced and enzymatically digested in RPMI as described above to obtain single cell suspension for downstream phenotyping. For that, cells were incubated with saturating concentrations of human immunoglobulins for 20 mins in RT and specific cell populations were detected with primary antibodies directly conjugated with fluorescent marker as listed in Table 2 for 30 minutes at 4°C. Populations of interest were analysed or sorted using a BD LSRFortessa™ Cell Analyzer and BD Influx™ Cell Sorter, respectively (both BD Biosciences). The population of interest was gated according to its FSC/SSC criteria. The dead cell population was excluded using 7-AAD staining (BD Biosciences). Data were analysed with the FlowJo™ software (BD Biosciences).

**Table 2.** Antibodies used in the study.

| Target | Visualisation | Dilution | Reference |
| --- | --- | --- | --- |
| CD45 | Brilliant Violet 421™ anti-human | 1:40 | BioLegend (368521) |
| CD31 | Brilliant Violet 421™ anti-human | 1:40 | BioLegend (303123) |
| CD140b (PDGFRβ) | PE anti-human | 1:20 | BioLegend (323605) |
| CD146 (MUC18) | Alexa Fluor® 488 anti-human | 1:20 | BioLegend (361019) |
| COL15A1 | Immunohistochemistry | 1:100 | Atlas (HPA017915) |
| D2-40 (Mesothelin) | Immunohistochemistry | prediluted | CellMarque (760-4395) |
| Podoplanin | APC/Cyanine7 anti-human | 1:100 | BioLegend (337029) |

### Quantitative real-time PCR (qPCR)

Total RNA was extracted using Qiagen RNeasy Kit according to the manufacturer”s recommendations. All RNAs were reverse transcribed with MultiScribe™ Reverse Transcriptase (ThermoFisher Scientific), following the manufacturer”s instructions. All qPCR reactions were performed with a CFX96 Touch Real-Time PCR System from Bio-Rad and the SYBR® Green JumpStart™ *Taq* ReadyMix™ (Sigma). Experiments were performed in triplicates for each data point. Each sample was normalized based on its expression of actin gene using ^2ΔΔ^CT-method. Data were analysed by one-way ANOVA with Tukey”s multiple comparisons test. The primers pairs used for this study are listed in Table 3.

**Table 3.**
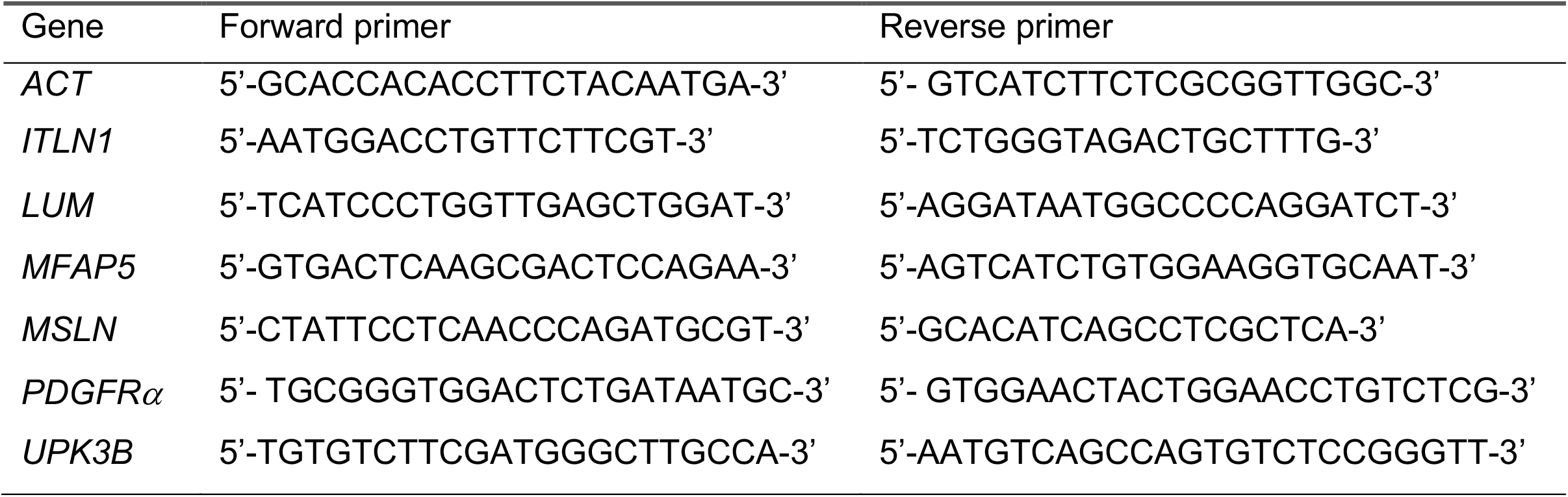
Primers used in the study.

### Electron microscopy

Low passage primary mesothelial cells were grown on Melinex plastic coverslips (TAAB) as described above. For imaging, cells were briefly rinsed in 0.9% saline and fixed in 2% glutaraldehyde/2% formaldehyde in 0.05 M sodium cacodylate buffer containing 2 mM CaCl2 at 4°C. Samples were very briefly dipped twice in de-ionised water to remove any buffer salts and quickly plunge-frozen by dipping into liquid nitrogen-cooled ethane. Then, samples were transferred to liquid nitrogen-cooled brass inserts and freeze-dried overnight in a liquid nitrogen-cooled turbo freeze-drier (Quorum K775X). Samples were mounted on aluminium SEM stubs using conductive carbon sticky pads (Agar Scientific) and coated with 25 nm gold using a Quorum K575X sputter coater. Samples were viewed using a FEI/ThermoFisher Scientific Verios 460 scanning electron microscope run at 2.0 kV and 50 pA probe current. Secondary electron images were acquired using either an Everhard-Thornley detector in field-free mode (low resolution) or a Through-Lens detector in full immersion mode (high resolution).

### Immunohistochemistry

*Formalin-fixed, paraffin-embedded pleura tissues sections were dewaxed and rehydrated, and antigen retrieval was done by heating the slides in 10 mM citric acid buffer (pH 6.0) for 15 min. Sections were incubated overnight at 4 °C in a humidified chamber with appropriate primary antibodies. All sections were counterstained with Gill”s haematoxylin and images were taken with Zeiss Axio Observer at 10x magnification*.

### Bulk RNA-seq and genomic data

Paired-end transcriptome reads were quality filtered and mapped to GRCh38 (ensemble build 98) using STAR-v2.5.0c^52^ with a standard set of parameters (https://github.com/cancerit/cgpRna). Resulting bam files were processed to get per gene read count data using HTSeq 0.7.2^53^ with parameters --stranded=reverse and -- mode=union. We calculated TPM (Transcripts per Million) values using the count and transcript length data for further downstream analysis. Mutations in BAP1 and NF2 were obtained from the Cell Model Passports https://cellmodelpassports.sanger.ac.uk.

## ACKNOWLEDGEMENTS

Funding: SJM was supported by the MRC (MCMB MR/V028669/1 and MR/R009120/1), EPSRC (EP/R03558X/1), Cambridge Biomedical Research Centre (BRC-1215-20014); British Lung Foundation (BLF), Asthma+Lung UK (ALUK), Royal Papworth Hospital, and the Victor Dahdaleh Foundation. MG and HEF were supported by the British Lung Foundation and Wellcome Trust grant 206194. RCR is supported by the Cambridge BRC, Cancer Research UK Cambridge Centre, BLF and Royal Papworth Hospital. NK was supported by the NIH R01HL141852, R01HL127349, U01HL145567, and U01HL122626.

## SUPPLEMENTARY LEGENDS

**Figure S1 scRNA-seq atlas of human parietal pleura by donor**

**A**. Uniform manifold approximation and projection (UMAP) of 64,514 pleural cells from 8 donors are colour-coded by donor.

**B**. Table of relative abundance by cell type and donor. Values are proportion of the 64,514 pleural cells presented as a percentage of the whole.

**Figure S2 Cell type identification**

UMAP projection of canonical marker gene expression: mesothelial cells (UPK3B, KRT18, KRT19, CALB2); fibroblasts (PDGFRA, LUM); smooth muscle cells (ACTA1); endothelial cells (CDH5, CLDN5, ERG, PODXL, TIE1); immune subtypes (CD45, CD2, CD4, CD79A).

**Figure S3 Patient contribution to each cell type group**

**A**. Relative abundance of stromal cell type and donor expressed graphically. Values are proportion of the 32,750 stromal cells.

**B**. Relative abundance by cell type and donor. Values are proportion of the 32,750 stromal cells presented as a percentage of the whole.

**Figure S4 Distribution of universal fibroblast markers**

**A**. Violin plot of *PI16*+ *COL15A1-* fibroblast marker gene expression.

**B**. Violin plot of *PI16*-*COL15A1+* fibroblast marker gene expression.

**Figure S5 Additional ANGPTL7+ fibroblast marker genes**

**A**. Violin plot of gene expression in ANGPTL7 fibroblasts

**B**. UMAP illustrating ANGPTL7+ cells.

**Figure S6 Stemness gene expression**

Violin plot of additional stemness-associated gene expression.

**Figure S7 Transcription factor expression in pleural fibroblasts**

**A**. Violin plot of selected transcription factor expression in pleural stromal cell populations

**B**. UMAP illustrating HES4+ cells.

**C**. UMAP illustrating ZNF281+ cells.

**Figure S8 Further myofibroblast markers**

Violin plot of myofibroblast marker gene (TAGLN, MYH1) expression.

**Figure S9 Further iCAF markers**

Violin plot of selected iCAF marker gene expression (AGTR1, HAS1, HAS2).

**Figure S10 Bulk RNA-seq analysis of healthy donor and mesothelioma cell cultures**

**A**. Gene Ontology (GO) Term analysis of mesothelioma cell bulk RNA-seq relative to bulk RNA-seq of healthy mesothelial cell cultures.

**B**. Similarly, KEGG pathway analysis of mesothelioma cell bulk RNA-seq relative to bulk RNA-seq of healthy mesothelial cell cultures.

**C**. Volcano plot of mesothelioma cell bulkRNA-seq relative to bulk RNA-seq of healthy mesothelial cell cultures (genes expressed more highly

