## Supplementary data for "Single-cell transcriptomic analysis of human pleura reveals stromal heterogeneity and informs in vitro models of mesothelioma"

FIGURE S1

A

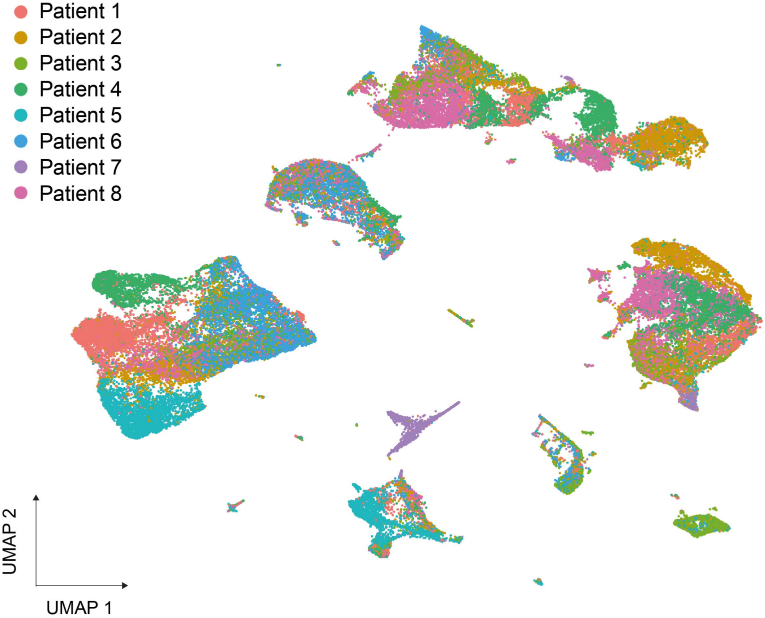


B

| | Mesothelial | Fibroblast | Endothelial | Smooth Muscle | Myeloid | T cells | B cells | Pericytes | Mast cells | Dendritic cells |
|---|---|---|---|---|---|---|---|---|---|---|
| Patient 1 | 2.00% | 2.900% | 6.00% | 0.60% | 0.60% | 0.20% | 0.00% | 0.10% | 0.10% | 0.00% |
| Patient 2 | 4.50% | 5.100% | 3.10% | 0.60% | 0.40% | 0.40% | 0.00% | 0.30% | 0.00% | 0.00% |
| Patient 3 | 1.90% | 3.000% | 2.80% | 0.90% | 0.40% | 1.30% | 1.80% | 0.30% | 0.00% | 0.10% |
| Patient 4 | 4.50% | 4.800% | 3.50% | 0.50% | 0.30% | 0.00% | 0.00% | 0.30% | 0.00% | 0.00% |
| Patient 5 | 0.40% | 0.600% | 5.30% | 0.10% | 3.10% | 0.10% | 0.00% | 0.10% | 0.20% | 0.20% |
| Patient 6 | 0.00% | 1.600% | 6.90% | 2.70% | 0.30% | 1.20% | 0.30% | 0.50% | 0.00% | 0.00% |
| Patient 7 | 0.70% | 0.400% | 0.00% | 0.00% | 0.10% | 2.30% | 0.00% | 0.00% | 0.00% | 0.00% |
| Patient 8 | 6.10% | 7.800% | 2.60% | 1.60% | 0.50% | 0.10% | 0.00% | 0.40% | 0.10% | 0.00% |

FIGURE S2

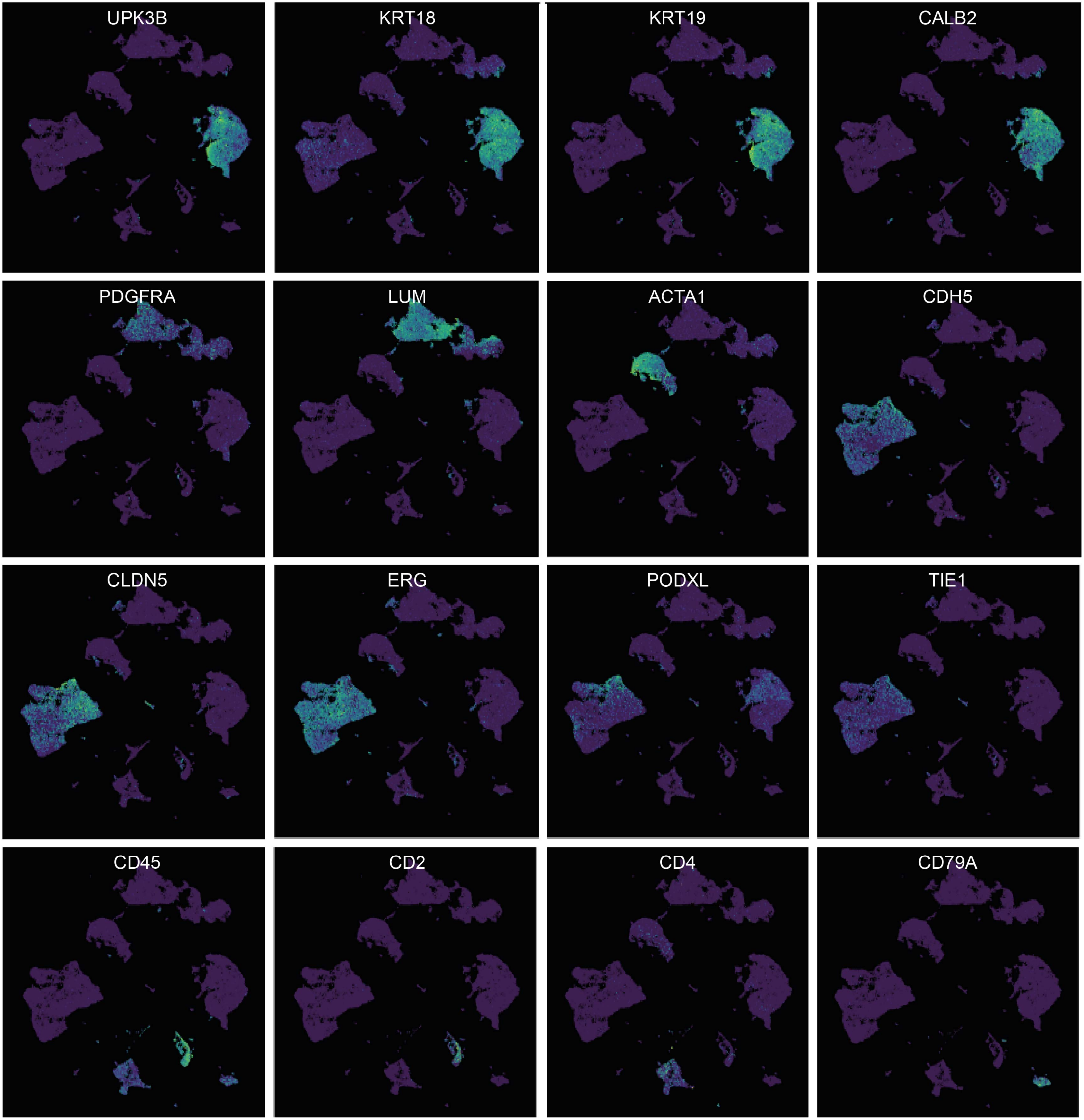
UPK3B
KRT18
KRT19
CALB2
PDGFRA
LUM
ACTA1
CDH5
CLDN5
ERG
PODXL
TIE1
CD45
CD2
CD4
CD79A

A

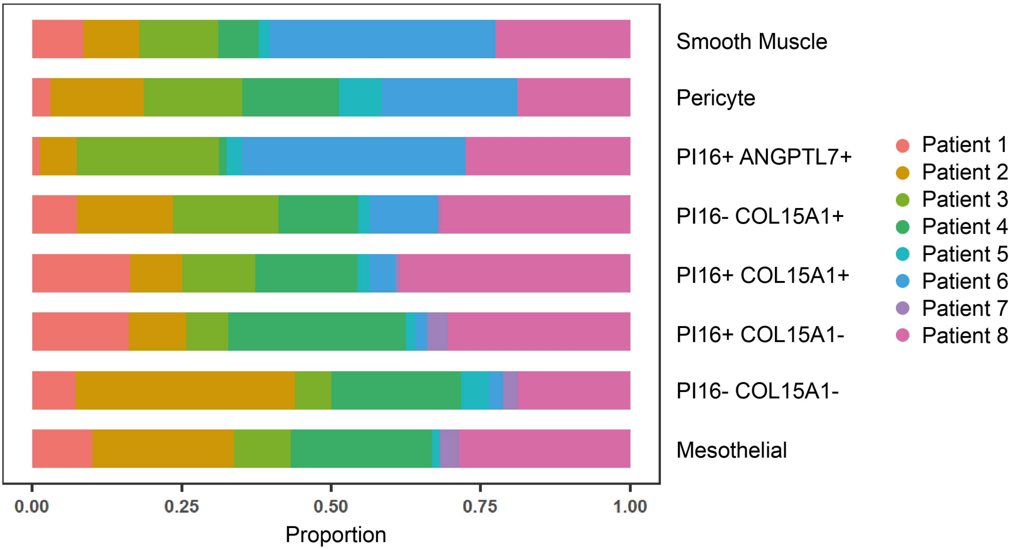
Smooth Muscle
Pericyte
PI16+ ANGPTL7+
PI16- COL15A1+
PI16+ COL15A1+
PI16+ COL15A1-
PI16- COL15A1-
Mesothelial
0.00
0.25
0.50
0.75
1.00
Proportion
Patient 1
Patient 2
Patient 3
Patient 4
Patient 5
Patient 6
Patient 7
Patient 8

B

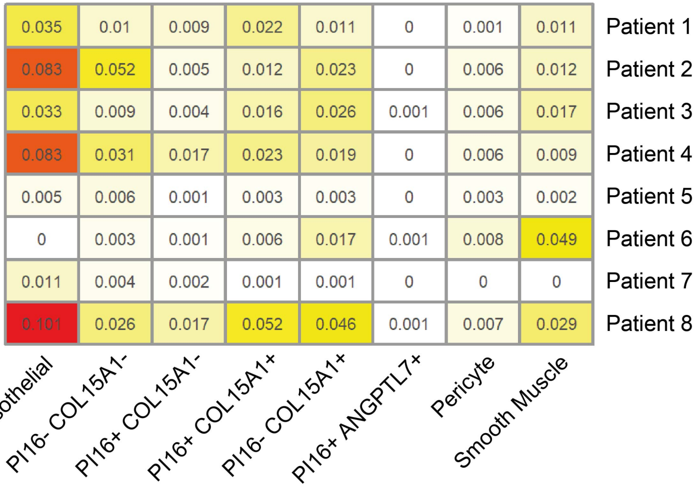
0.035 0.01 0.009 0.022 0.011 0 0.001 0.011 Patient 1
0.083 0.052 0.005 0.012 0.023 0 0.006 0.012 Patient 2
0.033 0.009 0.004 0.016 0.026 0.001 0.006 0.017 Patient 3
0.083 0.031 0.017 0.023 0.019 0 0.006 0.009 Patient 4
0.005 0.006 0.001 0.003 0.003 0 0.003 0.002 Patient 5
0 0.003 0.001 0.006 0.017 0.001 0.008 0.049 Patient 6
0.011 0.004 0.002 0.001 0.001 0 0 0 Patient 7
0.101 0.026 0.017 0.052 0.046 0.001 0.007 0.029 Patient 8
Mesothelial
PI16- COL15A1-
PI16+ COL15A1-
PI16+ COL15A1+
PI16- COL15A1+
PI16+ ANGPTL7+
Pericyte
Smooth Muscle

FIGURE S4

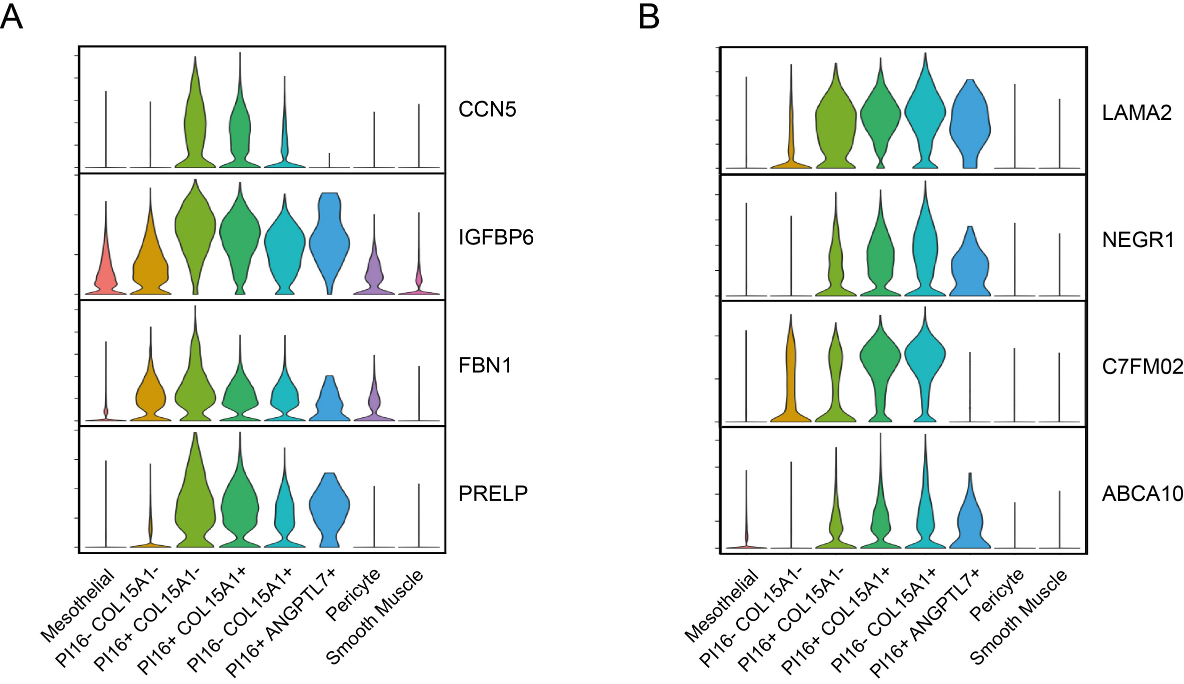
A
CCN5
IGFBP6
FBN1
PRELP
Mesothelial
PI16- COL15A1-
PI16+ COL15A1-
PI16+ COL15A1+
PI16- COL15A1+
PI16+ ANGPTL7+
Pericyte
Smooth Muscle
B
LAMA2
NEGR1
C7FM02
ABCA10
Mesothelial
PI16- COL15A1-
PI16+ COL15A1-
PI16+ COL15A1+
PI16- COL15A1+
PI16+ ANGPTL7+
Pericyte
Smooth Muscle

FIGURE S5

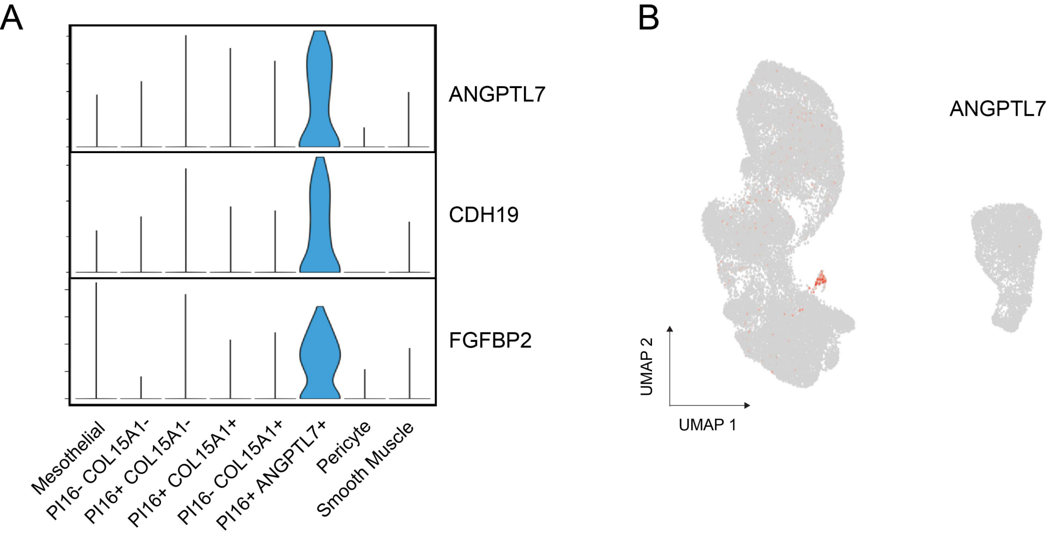
A
ANGPTL7
CDH19
FGFBP2
Mesothelial
PI16- COL15A1-
PI16+ COL15A1-
PI16+ COL15A1+
PI16- COL15A1+
PI16+ ANGPTL7+
Pericyte
Smooth Muscle
B
ANGPTL7
UMAP 2
UMAP 1

### FIGURE S6

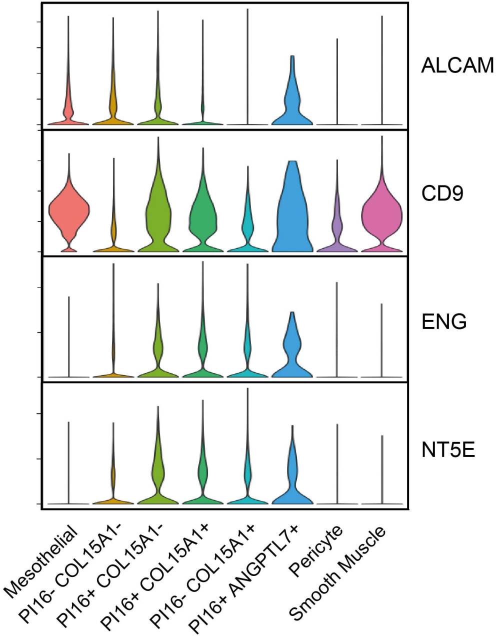
ALCAM
CD9
ENG
NT5E
Mesothelial
PI16- COL15A1-
PI16+ COL15A1-
PI16+ COL15A1+
PI16- COL15A1+
PI16+ ANGPTL7+
Pericyte
Smooth Muscle

FIGURE S7

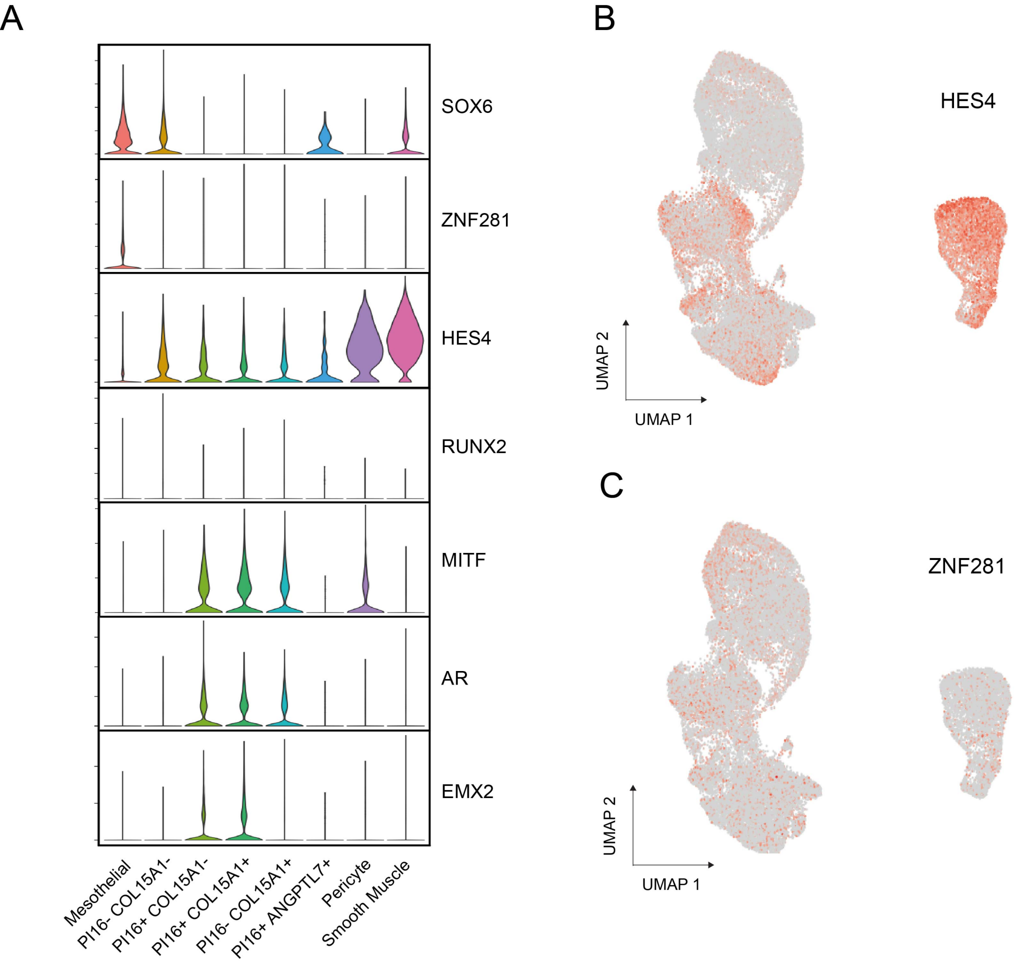

A
SOX6
ZNF281
HES4
RUNX2
MITF
AR
EMX2
Mesothelial
PI16- COL15A1-
PI16+ COL15A1-
PI16+ COL15A1+
PI16- COL15A1+
PI16+ ANGPTL7+
Pericyte
Smooth Muscle
B
HES4
UMAP 2
UMAP 1
C
ZNF281
UMAP 2
UMAP 1

### FIGURE S8

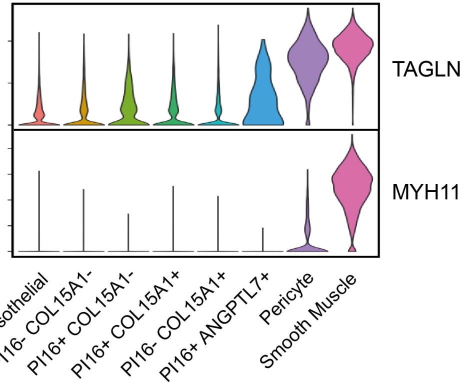

TAGLN
MYH11
Mesothelial
PI16- COL15A1-
PI16+ COL15A1-
PI16+ COL15A1+
PI16- COL15A1+
PI16+ ANGPTL7+
Pericyte
Smooth Muscle

### FIGURE S9

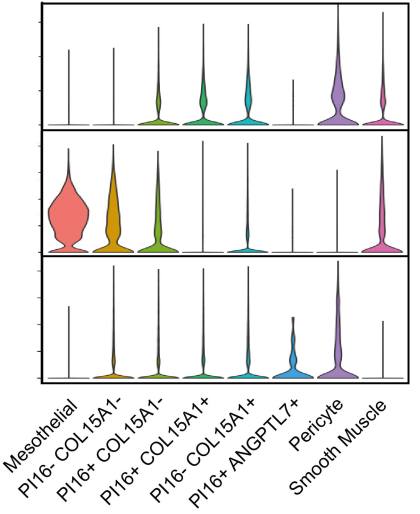

AGTR1
HAS1
HAS2
Mesothelial
PI16- COL15A1-
PI16+ COL15A1-
PI16+ COL15A1+
PI16- COL15A1+
PI16+ ANGPTL7+
Pericyte
Smooth Muscle

FIGURE S10

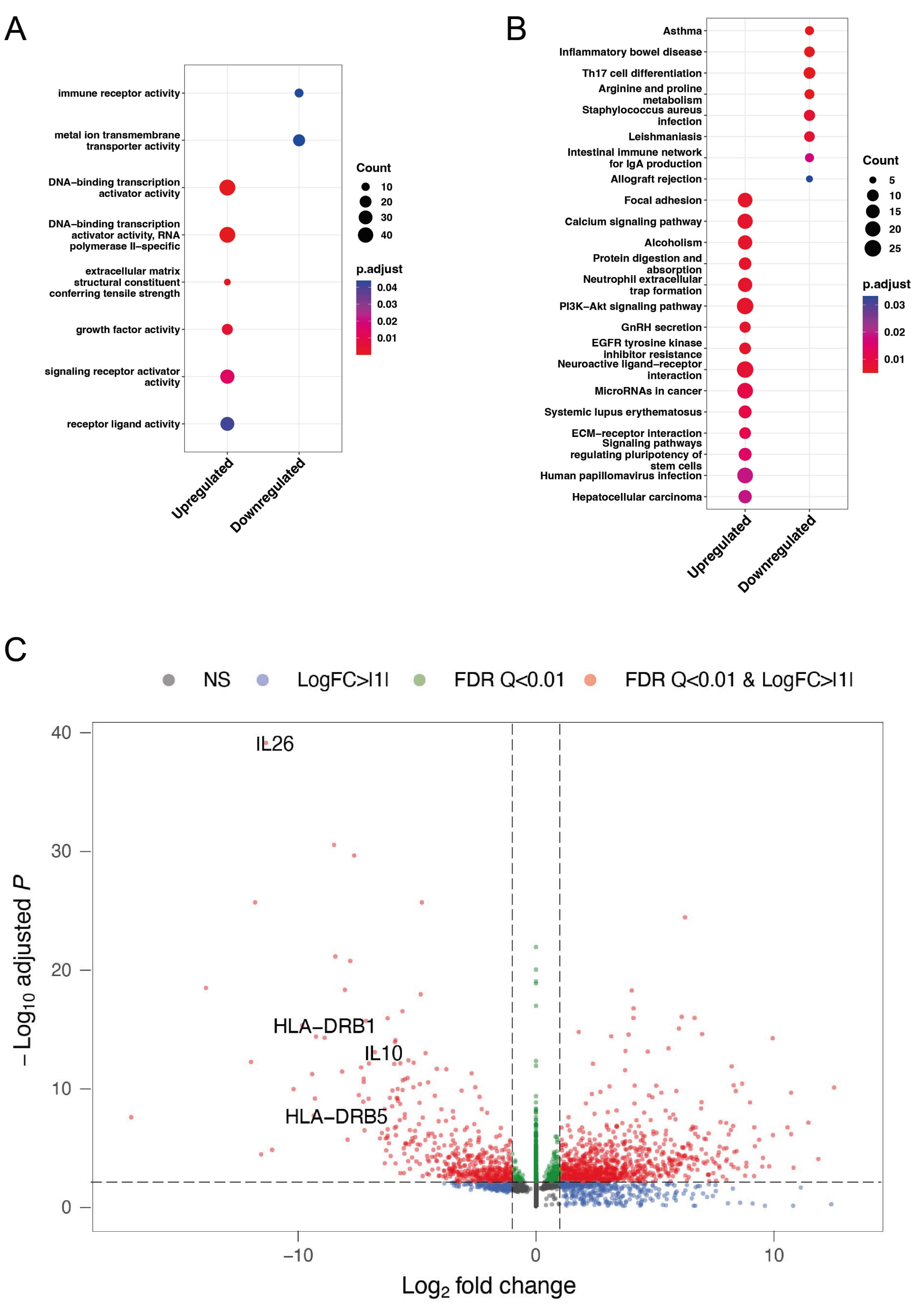

A
immune receptor activity
metal ion transmembrane transporter activity
DNA–binding transcription activator activity
DNA–binding transcription activator activity, RNA polymerase II–specific
extracellular matrix structural constituent conferring tensile strength
growth factor activity
signaling receptor activator activity
receptor ligand activity
Upregulated
Downregulated
Count
10
20
30
40
p.adjust
0.04
0.03
0.02
0.01
B
Asthma
Inflammatory bowel disease
Th17 cell differentiation
Arginine and proline metabolism
Staphylococcus aureus infection
Leishmaniasis
Intestinal immune network for IgA production
Allograft rejection
Focal adhesion
Calcium signaling pathway
Alcoholism
Protein digestion and absorption
Neutrophil extracellular trap formation
PI3K–Akt signaling pathway
GnRH secretion
EGFR tyrosine kinase inhibitor resistance
Neuroactive ligand–receptor interaction
MicroRNAs in cancer
Systemic lupus erythematosus
ECM–receptor interaction
Signaling pathways regulating pluripotency of stem cells
Human papillomavirus infection
Hepatocellular carcinoma
Upregulated
Downregulated
Count
5
10
15
20
25
p.adjust
0.03
0.02
0.01
C
NS
LogFC>I1I
FDR Q<0.01
FDR Q<0.01 & LogFC>I1I
IL26
HLA–DRB1
IL10
HLA–DRB5
40
30
20
10
0
−10
0
10
– Log10 adjusted P
Log2 fold change
